## Supplementary material for "Blocking toxin function and modulating the gut microbiota: caffeic acid phenethyl ester as a potential treatment for *Clostridioides difficile* infection"

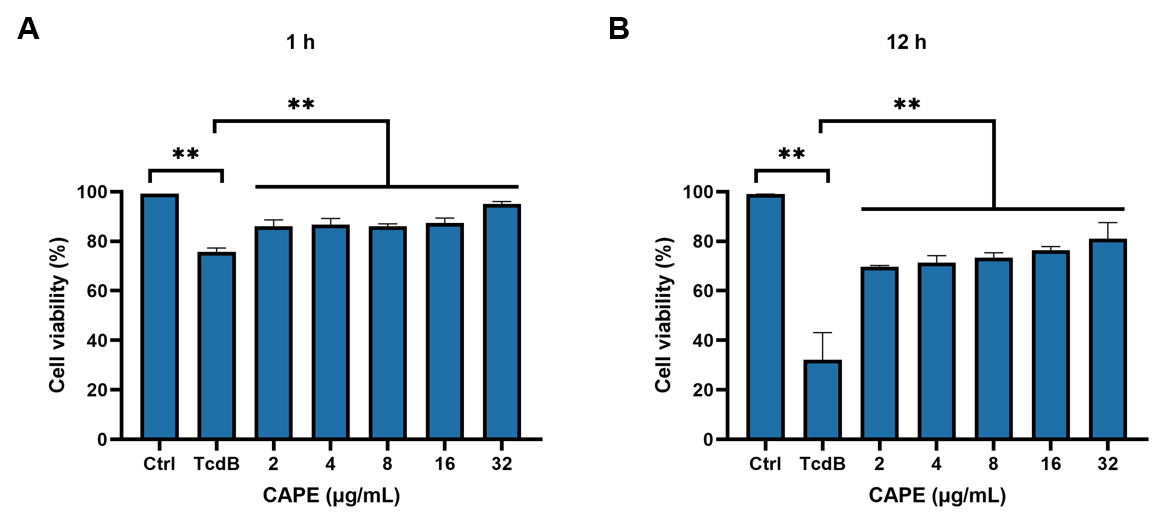

**Figure S1. CAPE reduces cell death induced by TcdB.**

TcdB was pretreated with CAPE for 1 h, followed by its addition to Vero cells for a 1 h (**A**) or 12 h (**B**) incubation to assess cell viability. * *p* <0.05; ** *p* <0.01 using one-way ANOVA.

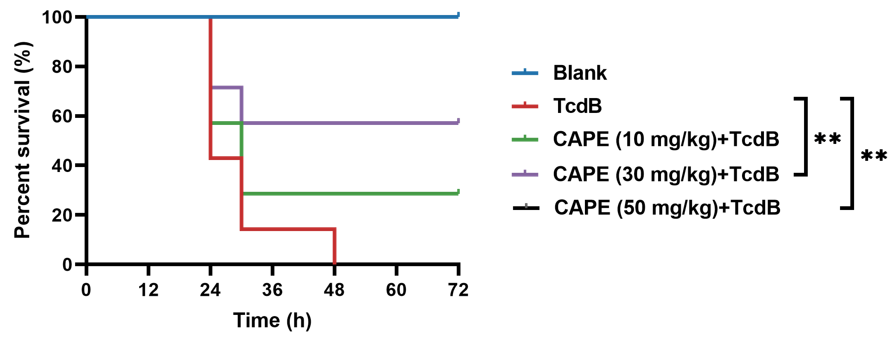
**Figure S2. CAPE neutralizes TcdB-mediated acute toxicity in mice.**

The survival rates of mice that were intraperitoneally injected with CAPE 2 h prior to the subsequent injection of native TcdB (30 ng). * *p* <0.05 and ** *p* <0.01 by log-rank (Mantel‒Cox) test.

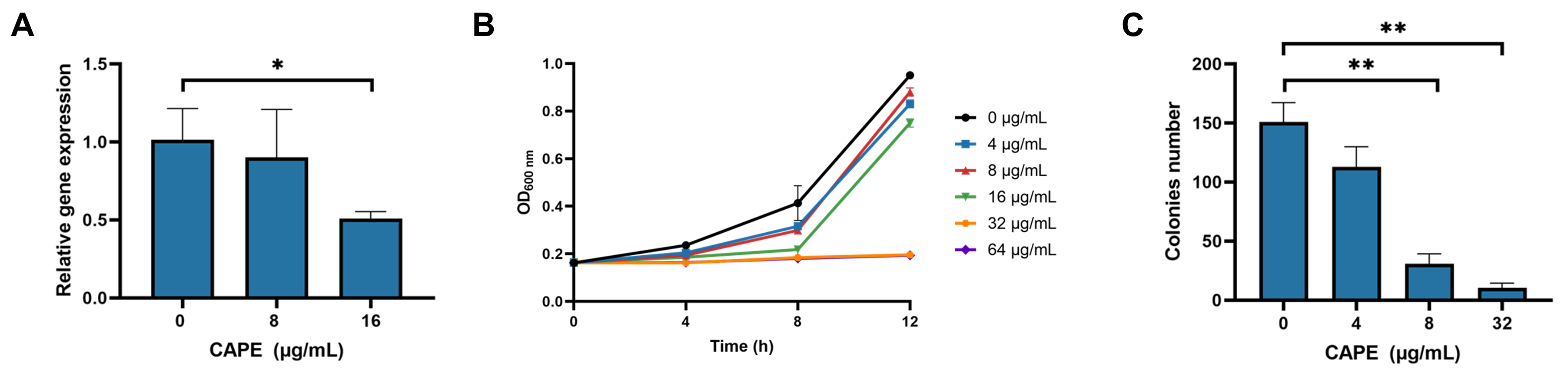

**Figure S3. CAPE inhibits the growth of *C. difficile* BAA-1870 and the expression of TcdB.**

(**A**) CAPE effectively suppresses the expression of TcdB in a dose-dependent manner. * *p* <0.05; ** *p* <0.01 using one-way ANOVA. (**B**) Growth curves of *C. difficile* BAA-1870 in the presence of varying concentrations of CAPE (0-64 µg/mL). (**C**) Effect of CAPE on spore production capacity of *C. difficile* BAA-1870. The colony forming units of heat-shocked spore solution of BAA-1870 progressively decline with the elevation of CAPE concentrations (0-32 µg/mL). * *p* <0.05; ** *p* <0.01 using one-way ANOVA.

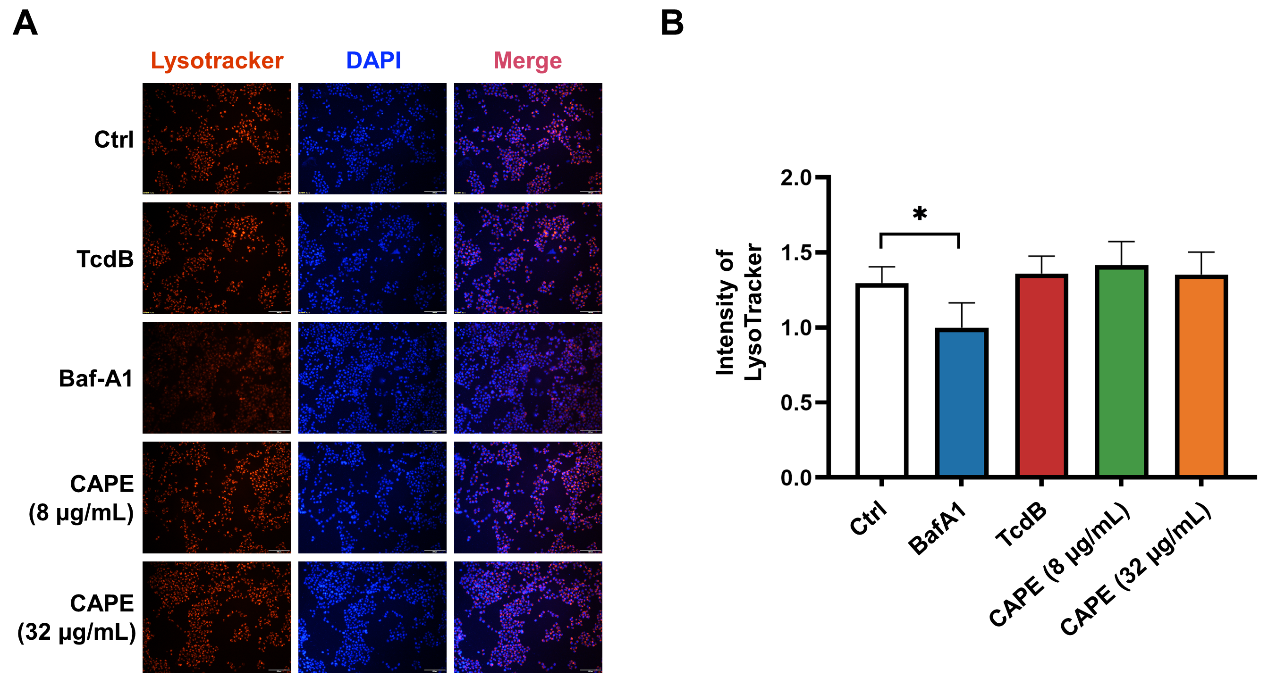

**Figure S4. Lysosomal activity of Caco-2 cells in the presence of CAPE.**

**(A)** Caco-2 cells were treated with 8 μg/mL or 16 μg/mL CAPE for 1 h. The acidic compartments were stained with LysoTracker Red DND-99, and the nuclei were stained with DAPI. Bar, 100 μm. **(B)** The intensities of LysoTracker staining were quantified using ImageJ. * *p* <0.05; ** *p* <0.01 using one-way ANOVA.

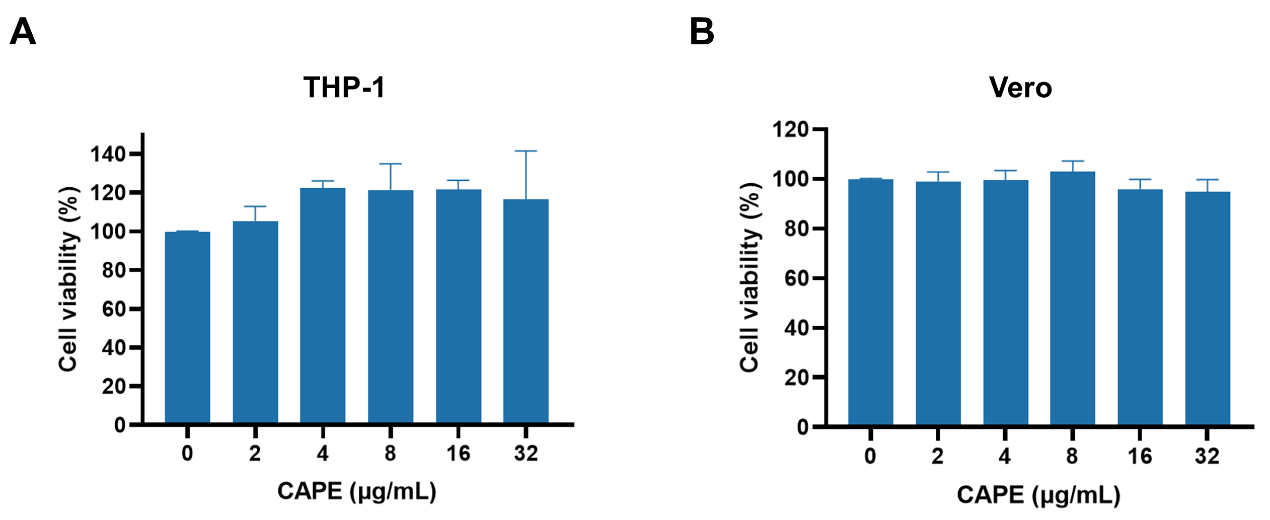

**Figure S5. CAPE is not cytotoxic.**

**(A-B)** THP-1 (**A**) and Vero (**B**) cells were incubated with increasing concentrations of CAPE for 3 h. Cytotoxicity was measured using a CCK-8 kit.

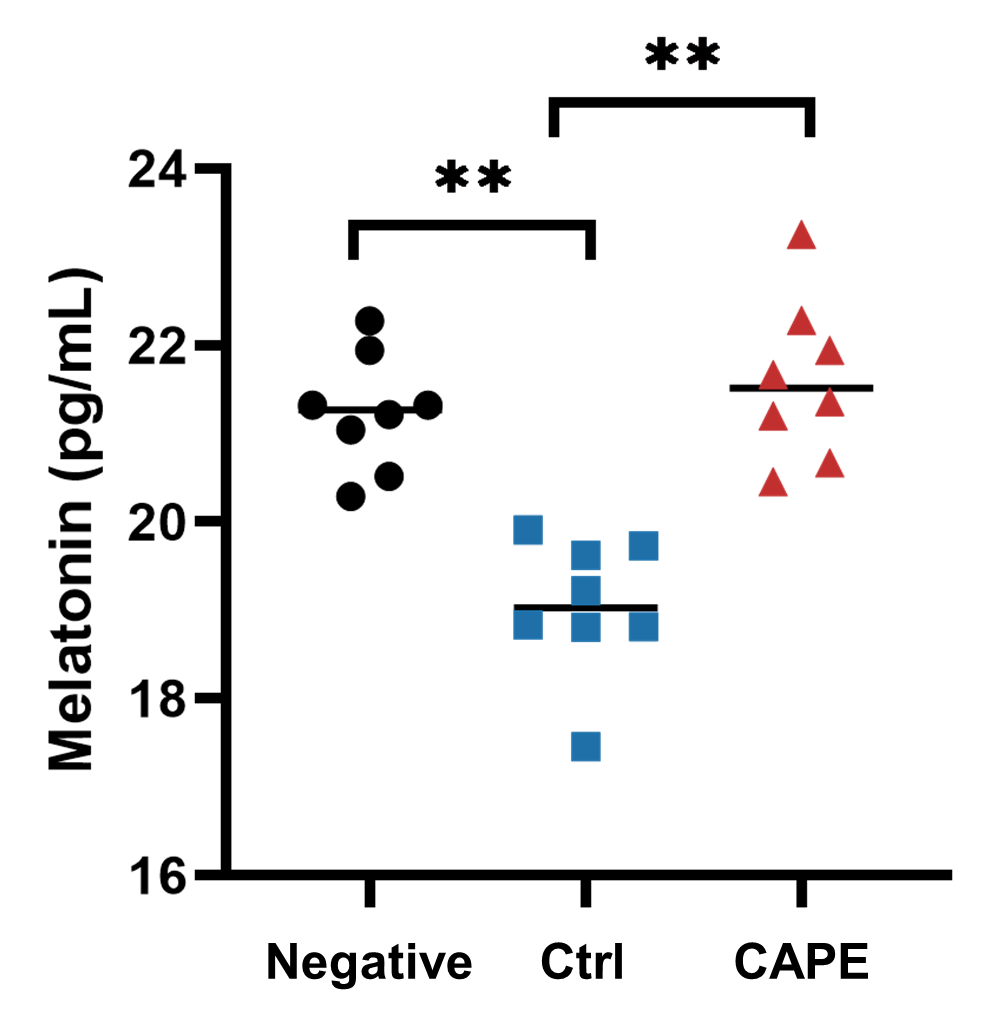

**Figure S6. Measurement of melatonin levels in mouse fecal samples.**

The melatonin content in mouse fecal supernatant was determined using a Mouse MT (melatonin) ELISA Kit. * *p* <0.05; ** *p* <0.01 using one-way ANOVA.

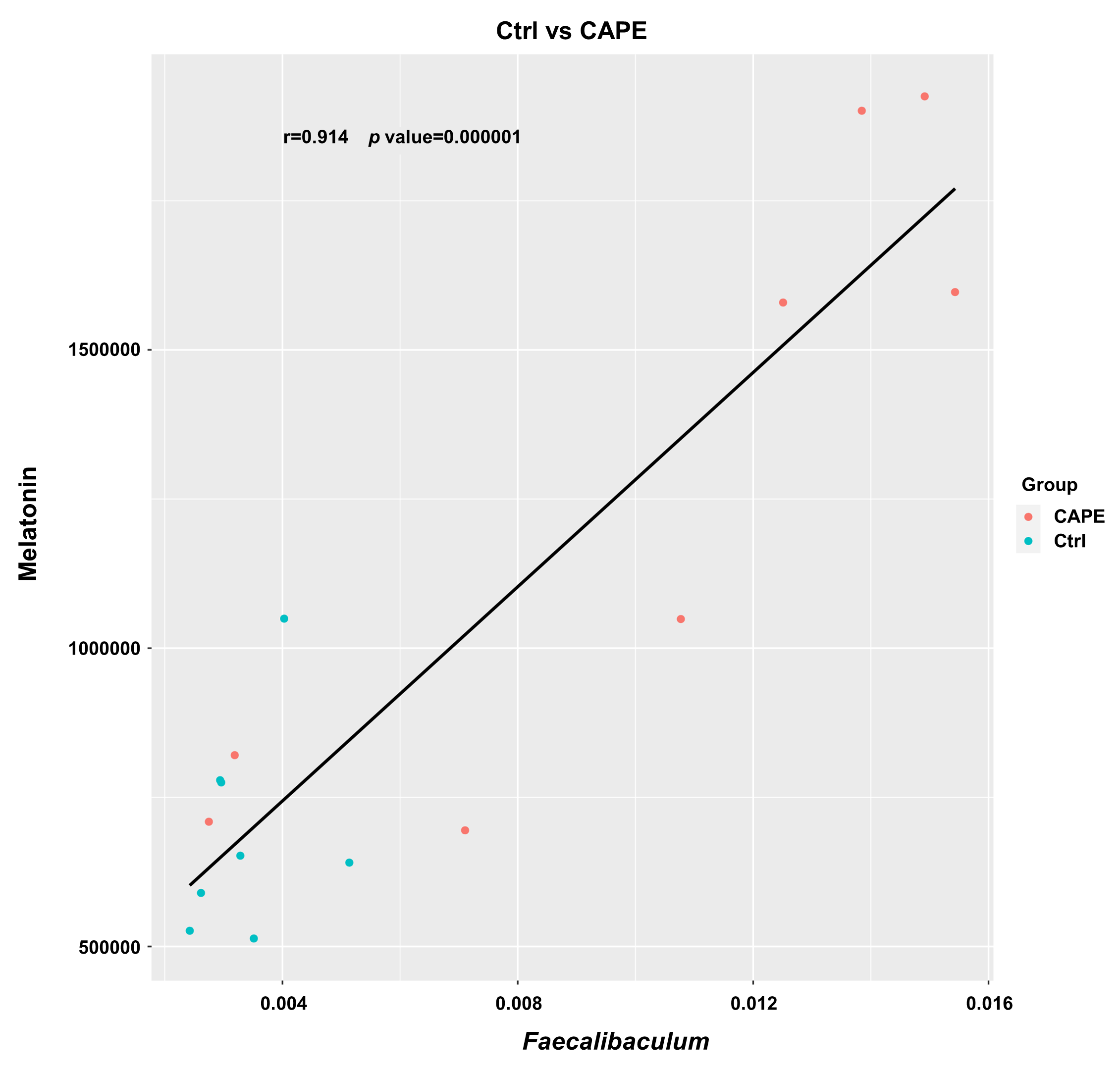

**Figure S7. The abundance of melatonin was plotted against the abundance of *Faecalibaculum* based on the microbiota and metabolome profiles.**

Correlation and statistical regression analyses were performed using Pearson’s method.

**Table S1.** **IC_50_ of the caffeic acid and its derivate on TcdB-mediated cell-rounding.**

| **Compound** | **Structure** | **IC_50_ (μg/mL)** |
| --- | --- | --- |
| Caffeic acid | 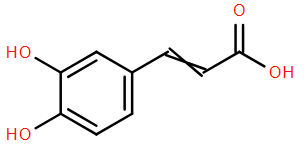 | 527.5 |
| Ethyl caffeate | 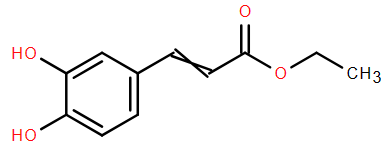 | 5.9 |
| CAPE | 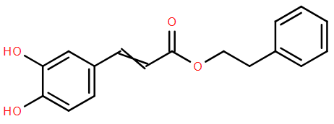 | 3.0 |
| Echinacoside | 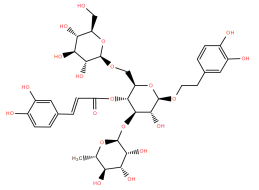 | 5.1 |
| Rosmarinic acid | 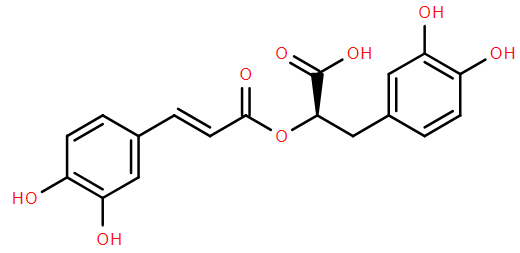 | 198.1 |
| Salvianolic acid | 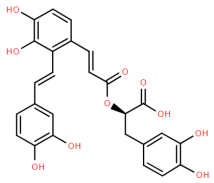 | 3.4 |
| Verbascoside | 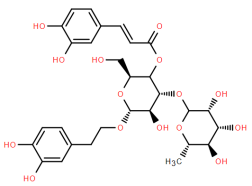 | 5.4 |

**Table S2. Primers used in this study.**

| **Primers** | **Sequences** | **Notes** | |
| --- | --- | --- | --- |
| YG101 | ctgGGATCCATGAGTTTAGTTAATAG | *gtd* BamHI-F | |
| YG102 | ctgGTCGACCTAAAGAGAACCTTCAA | | *gtd* SalI-R |
| YG103 | CAGAAGCAGCAGCTGCATTCCACCTTTCTACCAACTCTTGTTC | | *gtd*_L265A_ -F |
| YG104 | GAACAAGAGTTGGTAGAAAGGTGGAATGCAGCTGCTGCTTCTG | | *gtd*_L265A_ -R |
| YG105 | GTATTCCTGGTAACATATCAACAGCTAAATACATACCACCAATTTCT | | *gtd*_D286A_ -F |
| YG106 | AGAAATTGGTGGTATGTATTTAGCTGTTGATATGTTACCAGGAATAC | | *gtd*_D286A_ -R |
| YG107 | CAGGGCCACTTAAGTTAATAGCAGTTTTAACATCTGGGAAGAA | | *gtd*_T465A_-F |
| YG108 | TTCTTCCCAGATGTTAAAACTGCTATTAACTTAAGTGGCCCTG | | *gtd*_T465A_-R |
| YG109 | TGCTTCAGGGCCACTTAAGTTAGCAGTAGTTTTAACATCTGGGAAG | | *gtd*_I466A_-F |
| YG110 | CTTCCCAGATGTTAAAACTACTGCTAACTTAAGTGGCCCTGAAGCA | | *gtd*_I466A_-R |
| YG111 | CTTGCATCGTCAAATGACCATGCGCTAGCCATTTCTTGTTCAGT | | *gtd*_L519_ -F |
| YG112 | ACTGAACAAGAAATGGCTAGCGCATGGTCATTTGACGATGCAAG | | *gtd*_L519A_ -R |
| YG113 | CTCTTGCATCGTCAAATGACGCTAAGCTAGCCATTTCTTGTTC | | *gtd*_W520A_ -F |
| YG114 | GAACAAGAAATGGCTAGCTTAGCGTCATTTGACGATGCAAGAG | | *gtd*_W520A_-R |

**Table S3. Sequence of primers used for Reverse transcriptase PCR (RT-PCR) analysis.**

| **Gene** | **Primer** | **Sequence (5’-3’)** |
| --- | --- | --- |
| *tcdB* | Forward | TAGCAAGAAGTTCAGAGAGAGGAT |
|  | Reverse | CTTGTATTTCACCATCTCCAGGA |
| *16S* | Forward | GCGTAGGCGGTCTTTCAAG |
|  | Reverse | TCGCAACTGGTGTTCCTCC |
